# Transcriptomic perspective on acromegaly- related PIT-1/SF-1 positive pituitary tumors

**DOI:** 10.1101/2025.05.28.656330

**Authors:** Julia Rymuza, Qilin Zhang, Mateusz Bujko

## Abstract

Neuroendocrine pituitary tumors (PitNETs) are classified based on clinical manifestation and expression of pituitary cell lineage-specific transcription factors (TFs) and hormones. A subtype of acromegaly-related tumors was found to express PIT-1 and SF-1 TFs, two markers of distinct pituitary cells. They were considered as multilineage or “somatogonadotoph” tumors. The aim of our study was to determine their identity and cell type origin by extensive transcriptomic analysis. For this purpose, we analyzed the RNA sequencing (RNAseq) data from 546 PitNETs (including 193 acromegaly-related tumors) and single cell RNAseq data from somatotroph and gonadotroph tumors and normal pituitary. Somatrotroph PitNETs co-expressing PIT-1 and SF-1 TFs were found in each included RNAseq dataset. Their transcriptomic profile and activity of pituitary TFs closely resembles other somatotroph tumors, but they substantially differ from gonadotroph PitNETs yet retain activity of *NR5A1* (SF-1) and expression of some of SF-1-regulated genes (*LHB* and *GNHRH*). SF-1 apparently regulates slightly different set of genes in double positive somatotroph PitNETs and gonadotroph tumors. Based on scRNAseq data a subcluster of normal gonadotroph cells with *POU1F1* (PIT-1) expression was identified, but tumor cells of PIT-1/SF-1 PitNETs were not similar to this subtype of normal gonadotrophs. A difference in genes expression profiles between 3 subtypes of somatotroph tumors was determined by analysis of both bulk- and scRNAseq data. From transcriptomic perspective of genes-coregulation and the activity of pituitary transcription factors, acromegaly-related PitNETs co-expressing PIT-1 and SF-1 are subtype of PIT-1 lineage tumors and molecular data do not support considering them as multilineage.

## Introduction

Neuroendocrine pituitary tumors (PitNETs) also named pituitary adenomas are among the most common intracranial tumors in adults. They are a heterogenous group of tumors, that originates from distinct pituitary secretory cells. Pituitary tumors are classified according to the recommendations of fifth edition of Classification of Endocrine and Neuroendocrine Tumors [1]. The principle of this classification is the paradigm that mature pituitary cell that secrete specified tropic hormones originate from common progenitor cells through the differentiation process that is orchestrated by a set of particular transcription factors. The key transcription factors for development of mature corticotroph and gonadotroph pituitary lineages are TPIT (encoded by *TBX19* gene) and SF-1 (*NR5A1*), respectively. In turn, PIT-1 (*PUO1F1*) transcription factor was found pivotal in maturation of somatotroph, lactotroph and tyrotroph cells.

Thus, somatotroph pituitary tumors (sPitNETs) are by definition those that express PIT-1 transcription factor and produce growth hormone. In addition to somatotroph tumors, other subtypes of GH- and PIT-1-positive tumors cause acromegaly: mammosomatotroph tumors and mature plurihormonal tumors (characterized by expression of GH and additional hormones (prolactin (PRL) and thyrotropic hormone, respectively), mixed somatolactotroph tumors (consisting of two distinct populations of somatotroph and lactotroph tumor cells), or low-differentiated PIT-1 tumors [2,3].

According to a recently published data, somatotroph pituitary tumors that cause acromegaly fall into 3 distinct molecular subtypes [4–7]. They differ in gene expression, DNA methylation and DNA copy number profiles, as well as in the frequency of *GNAS* mutations, granularity scores on electron microscopy images, percentage of invasive tumors and tumor size [4–6,8]. By analyses of genes expression, we found that one of these three subtypes (referred as Subtype 1 in our previous report [7]) is characterized by the expression of *NR5A1* (SF-1), a marker of gonadotroph PitNETs. Such tumors with expression of both PIT-1 and SF-1 TFs were occasionally reported by the others through the years [9–15] and they were pointed as a matter of special interest. These PIT-1/SF-1 positives sPitNETs are different in many ways as compared to the other acromegaly-related PitNETs without SF-1 expression. They are densely granulated adenomas without *GNAS* mutation, with ectopic *GIPR* expression, distinguished DNA methylation profile and high level of genomic instability [4,6,7].

Recent discussions of PIT-1/SF-1 tumors have been marked by confusion, as these two TFs are the markers specific to separate pituitary linages according to official classification. These tumors were commonly referred to as multilineage pituitary tumors [16,17], gonado/somatotroph tumors [6], somatogonadotroph [18] or plurihormonal tumors [9,11]. In this article we made an attempt to reveal the identity of double positive PIT-1/SF-1 tumors by in depth analysis of combination of our in house generated and publically available transcriptomic data on pituitary tumors as well as the available data from scRNAseq experiments in PitNETs and normal pituitary. The main question we addressed are: does PIT-1/SF-1 are more related to gonadotrop or somatotroph PitNETs and does experimental data justifies considering them as multilineage; are there normal counterparts of double positive PIT-1/SF-1 cells in pituitary; what distinguishes PIT-1/SF-1 and other PIT-1 somatotroph tumors at genes expression level. We are convinced that transcriptomic point of view with a focus on the transcription factors activity is the perspective underlying official WHO classification of pituitary tumors.

## Methods

### Bulk RNASeq data

We used RNAseq data previously generated by our research groups HRA003588 [19] and E-MTAB-11889 [8]. Through the literature review we also identified additional 3 datasets that contain bulk transcriptomic data of PitNETs including E-MTAB-7768 [15], GSE213527 [20], GSE209903 [21]. Depending on available data, we used either Salmon [22] to map reads to human genome version hg38 and taximport [23] to extract row counts from row reads or featuresCounts [24] to count reads from already mapped reads. In the analysis, we used genes detected in all studies that had more than 10 reads in over 25% of samples. The dataset was then variance stabilized transformed (VST) using DESeq2 [25] and batch effects were removed with the removeBatchEffect function from the limma package [26]. Differentially expressed genes were detected using linear model based on transformed, batch corrected data with false discovery rate (FDR) p-value correction.

To identify previously established transcriptomic subtypes of sPitNETs in the analyzed cohort, we focused on samples which were diagnosed with acromegaly or mixed acromegaly and hyperprolactinemia or had serum IGF-1 level above 400 ng/mL. We clustered samples using k-means clustering, hierarchical clustering and Gaussian Mixture Model. We assign samples to a cluster by choosing the most abounded cluster based on three clustering methods. For samples with discordant results among algorithms, we assign them based on previously established marker genes for sPitNET subtypes [8].

To further investigate expression patterns, we used WGCNA [27] to find modules of highly correlated genes and relate them to chosen data features. We also computed regulons, which are groups of genes co-regulated by a specific transcription factor (TF) with the RTN package [28]. For TF with documented function in pituitary (list in Supplementary Table S1) we calculated their activity in the samples. We used two-tailed GSEA to check if DEGs are enriched with regulons [29,30].

### Single cell RNAseq data

We utilized the previously obtained scRNAseq data PRJCA009690 [31] as well as the other publicly available single cell transcriptomes of both adult pituitary gland (APG) and PitNETs: GSE208108 [32]; HRA003110 [33]; HRA003483 [34]; PRJNA1137596 [35] as well as fetal pituitary (FPG) GSE142653 [36]. Raw reads were processed with alvin-fry [37] through simpleleaf framework using GRCh38 genome assembly. All the data was further analyzed using Seurat package (v5.1.0) [38]. Low quality cells were filtered based on nFeature_RNA, nCount_RNA, percent.mt. Doublets were detected using scDblFinder [39] and only singlets were kept for further analysis. Red blood cells were identified by calculating RBC_score, with *HBB, HBA1, HBG2, HBA2* used as marker genes. We selected highly variable features, scaled and did principal component analysis (PCA) of the filtered count matrix. Depending on data complexity, we integrated the data using either CCA [40] or FastMNN [41] algorithms. Cluster cell types were identified based on expression of classical marker genes [31,32]. We assessed the similarity of tumors, using AddModuleScore with 20 top DEG specific to given clusters from APG.

We used Slingshot[42] and tradeSeq [43] to construct lineage trajectory of integrated APG and FPG gonadotroph cells and to examine gene expression with pseudotime. For integrated data from APG and somatotroph tumors, we use Monocle3[44] to calculate pseudotime-related trajectory.

Using decoupleR [45] we constructed pseudobulks of somatotroph cells in each sample. We applied pyDESeq2 [46] to calculate DEGs between somatotroph cells from each subtype and APG. For resulting gene sets, we conducted GSEA with decoupleR using KEGG database [47].

To compare regulons of important TF between somatotroph cells from Subtype 1 and gonadotrhop cells from gonadotrhop tumors, we used hdWGCNA [48]. We selected genes for each TF which met criteria of regulatory score above 0.75.

We inferred copy numbers of each sample with CopyKAT [49]. We used somatotroph cells from APG as normal cells. For each sample, we calculated CN profiles as average CN ration for chromosome arm. Each cell was then assigned to a stability cluster depending on aneuploidy score (AS) calculated as a number of chromosome arms with average CN ration above 0.1 after normalization by subtracting CN ration of chr22, to remedy previously showed shortcomings of only depth based CN calculations [4]. If AS was above 2, we designated cells as unstable, and if they had deletion of chr11 and less than 4 amplifications of chromosomal arms as del11.

## Results

### Clustering of pituitary tumor samples

Three molecular subtypes of somatotroph tumors causing acromegaly were identified in previous studies, therefore we started our analysis with an attempt of assigning each sample in study cohort comprising 193 acromegaly-causing tumors to one of these subtypes. Dimensionality reduction of transcriptomic data with UMAP showed a clustering of acromegaly-related tumors into three groups (Fig. 1A). Based on consensus of 3 clustering algorithms, 98% of the samples was assigned to one of these groups. Each group was identified as representing one of previously characterized subtypes (subtypes 1,2 and 3) [4,6,7] based on the presence of the samples from previous research with known transcriptome subtype assignment. Three samples (1.5% of all acromegaly-related PitNETs) with unclear annotation due to lack of consensus between 3 algorithms were assigned manually based on previously proposed marker genes [7] (Supplementary Fig. S1). Together, it allowed for the annotation of each acromegaly-related tumor sample in the study cohort with one of the known, previously described subtype of somatotroph tumors. Subtype 1, previously identified as composed of densely granulated SF-1/PIT-1 double positive tumors was represented by 43/193 (22%) of the samples; 83/193 (43%) samples were assigned to Subtype 2 previously identified as enriched for *GNAS*-mutated DG tumors; whereas 67/193 (35%) were of Subtype 3 known to be enriched for SG somatotroph PitNETs. According to previous observations, the co-expression of *POU1F1* and *NR5A1* was clearly confirmed in the tumors of Subtype 1 (Fig 1B). All these tumors were originally diagnosed as GH-omas or plurihirmonal tumors co expressing GH and PRL with one case of mixed GH/PRL and one of unknown diagnosis.

**Fig. 1.**
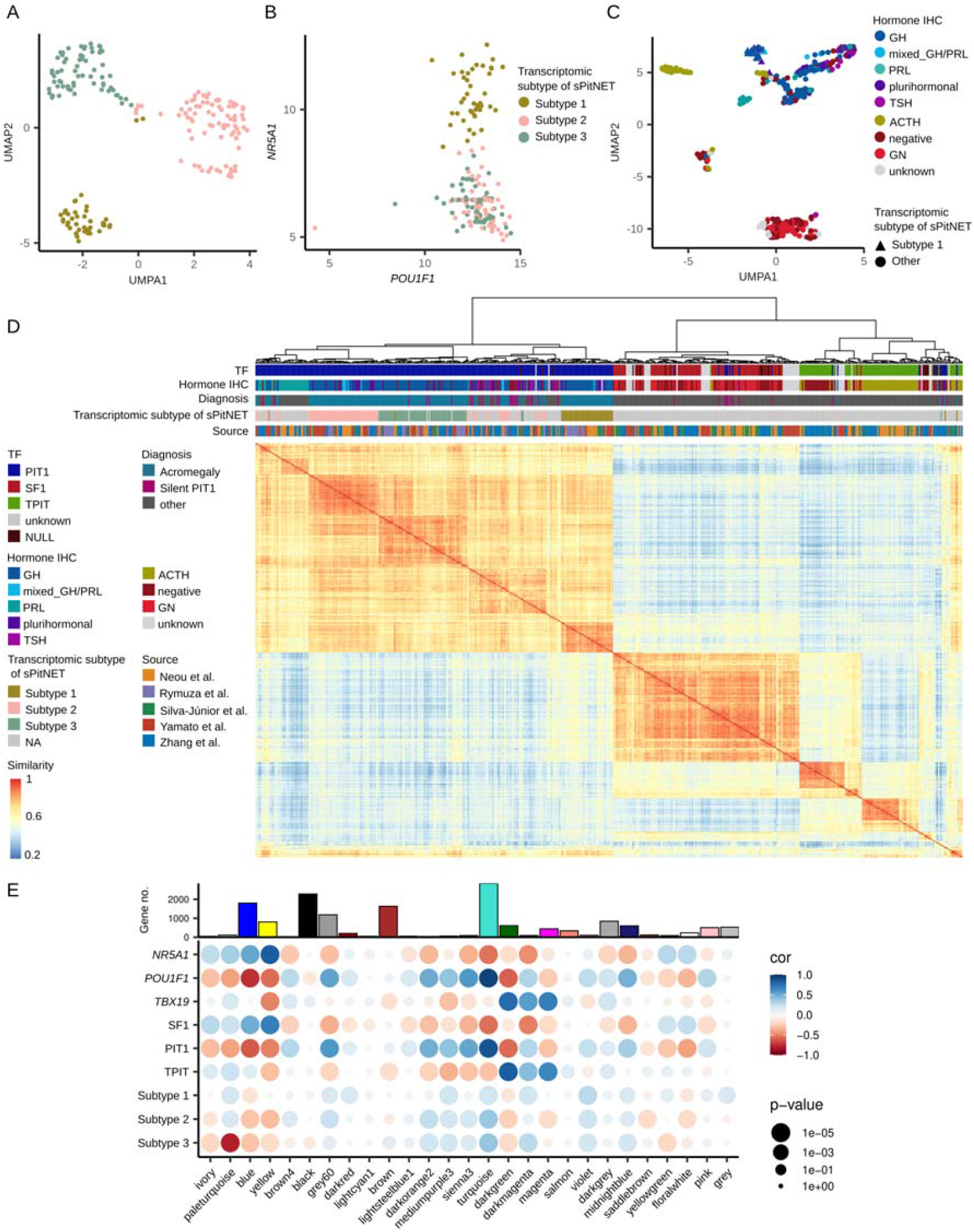
Transcriptomic profile of acromegaly-related pituitary tumors as well as all the PitNETs included in study cohort. A) UMAP of acromegaly-related tumors with assigned transcriptomic subtypes. B) Expression of *POU1F1* and *NR5A1* in acromegaly-related tumors samples colored according to transcriptomic subtype. C) UMAP of the whole PitNET study cohort labeled according to hormone staining by immunohistochemistry (IHC) with distinguished samples from acromegaly-related Subtype 1 tumors. D) Hierarchical clustering and similarity matrix of the transcriptomes of the entire study cohort with summary of molecular, histological and clinical characteristics. E) Results of Weighted Gene Co-expression Network Analysis (WGCNA) analysis, showing correlation (cor) between calculated gene modules and given features.

Having the acromegaly-related tumors characterized, we performed the unsupervised clustering of all the PitNETs in the study cohort (n=546) (Supplementary Table S2). We paid a special attention to the similarity of double positive *POU1F1*-and *NR5A1*-expressing tumors to the other histological types. The samples clustered according to the expression of lineage-specific TFs into PIT-1, TPIT and SF-1 tumors separately (Fig. 1C). Most of the tumors described as negative for TF expression (null cell tumors) (13/24) clustered with SF-1 group; 8/24 null cell tumors clustered with TPIT samples and 3/24 with PIT-1 cluster. The combined study cohort included 45 non-functioning PitNETs without reported immunohistochemical staining status i.e. no information on hormone nor transcription factor expression. These samples clustered mostly with SF-1 group (32 samples); 11 of them grouped with TPIT tumors and 2 with PIT-1. Importantly, all the acromegaly-related Subtype 1 tumors (positive for both *POU1F1* and *NR5A1*) form a separate cluster located closely to the PIT-1 but not to SF-1 tumors reflecting their much higher similarity to PIT-1 positive than gonadotroph PitNETs. This clustering pattern obtained with UMAP was confirmed with hierarchical clustering and similarity matrix (Fig. 1D). The dendrogram that represent the similarity of the samples contain the main 3 branches of PIT-1, SF-1 and TPIT tumors. With the exception of 3 samples, all double positive PIT-1/SF-1 tumors were grouped as a separate cluster within PIT-1 dendrogram branch. Of note, none of these tumors clustered within the same branch as gonadotroph SF-1-positive PitNETs.

### Genes co-expression analysis

Following the observation of much higher similarity of double positive PIT-1/SF-1 tumors to the other PIT-1 lineage tumor than SF-1 gonadotroph tumors, we looked whether the genes expression profile reflect their pituitary lineage identity. We used a Weighted Gene Co-expression Network Analysis (WGCNA) of the RNAseq data of the entire study cohort to generate gene co-expression network and to identify the groups of co-expressed genes. A total of 27 modules of co-expressed genes labeled with a distinct color names were found. We determined the relationship between each module and the expression of genes encoding for lineage specific transcription factor (*NR5A1, POU1F1* and *TBX19*). We also checked the relationship between modules and given subtypes of tumor sample including, SF-1 positive gonadotrophs and TPIT-positive corticotrophs and general PIT-1 lineage tumors (including those *POU1F1/NR5A1* expressing) as well as each of 3 transcriptomic subtypes of acromegaly-related tumors (Fig. 1E). We observed that the tumors of basic histological subtypes, defined by the expression of lineage-specific TF, have similar relationship with the co-expression modules as the expression of the gene encoding these respective TFs. PIT-1 tumors have similar relationship with WGCNA modules as *POU1F1* expression, while SF-1 gonadotroph tumors and TPIT corticotroph tumors have a similar relationship with the expression of *NR5A1* and *TBX19*, respectively. All three subtypes of acromegaly – related tumors were clearly most similar to PIT1-lineage tumors and *POU1F1* expression level in terms of the relationship with the WGCNA modules (Fig. 1E). Subtype 1 double positive tumors were similar to SF-1 - positive goadotropinomas in the relationship to only one module (grey).

In general, this unsupervised analysis of genes co-expression confirmed the observation from sample clustering, that the double positive Subtype 1 tumors are much more similar to other tumors of PIT1-lineage than that to SF-1-positive gonadotroph. There is a very little similarity in genes co-expression between PIT-1/SF-1- and SF-1-positive tumors.

### Regulon analysis

Transcriptomic data allows for the evaluation of the transcription factors activity. As the pituitary lineage-specific TF are substantial for official PitNETs classification and a number of TFs that orchestrate pituitary development were identified, we recognized regulon analysis as especially valuable.

Regulon (set of genes regulated by known transcription factor) was established for the entire dataset based on mutual information and subsequently the activity of all the known transcript factors was calculated for each tumor sample. Based on the literature data, we selected 25 TFs known to be relevant for differentiation and functioning of anterior pituitary cells and the regulons of 20 of these TFs were identified (5 TFs didn’t pass a default filtration due to small regulon size) (Supplementary Table S3). As the result, we established the regulons patterns characteristic for basic PitNETs histological subtypes with a distinct TFs activity in PIT-1 linage tumors, SF-1 gonadotropinomas and TPIT-corticotropinomas (Fig. 2A). Interestingly, the results suggest that each TF regulate mostly distinct set of genes with relatively low number of genes regulated by more than one TF (Fig. 2B).

**Fig. 2.**
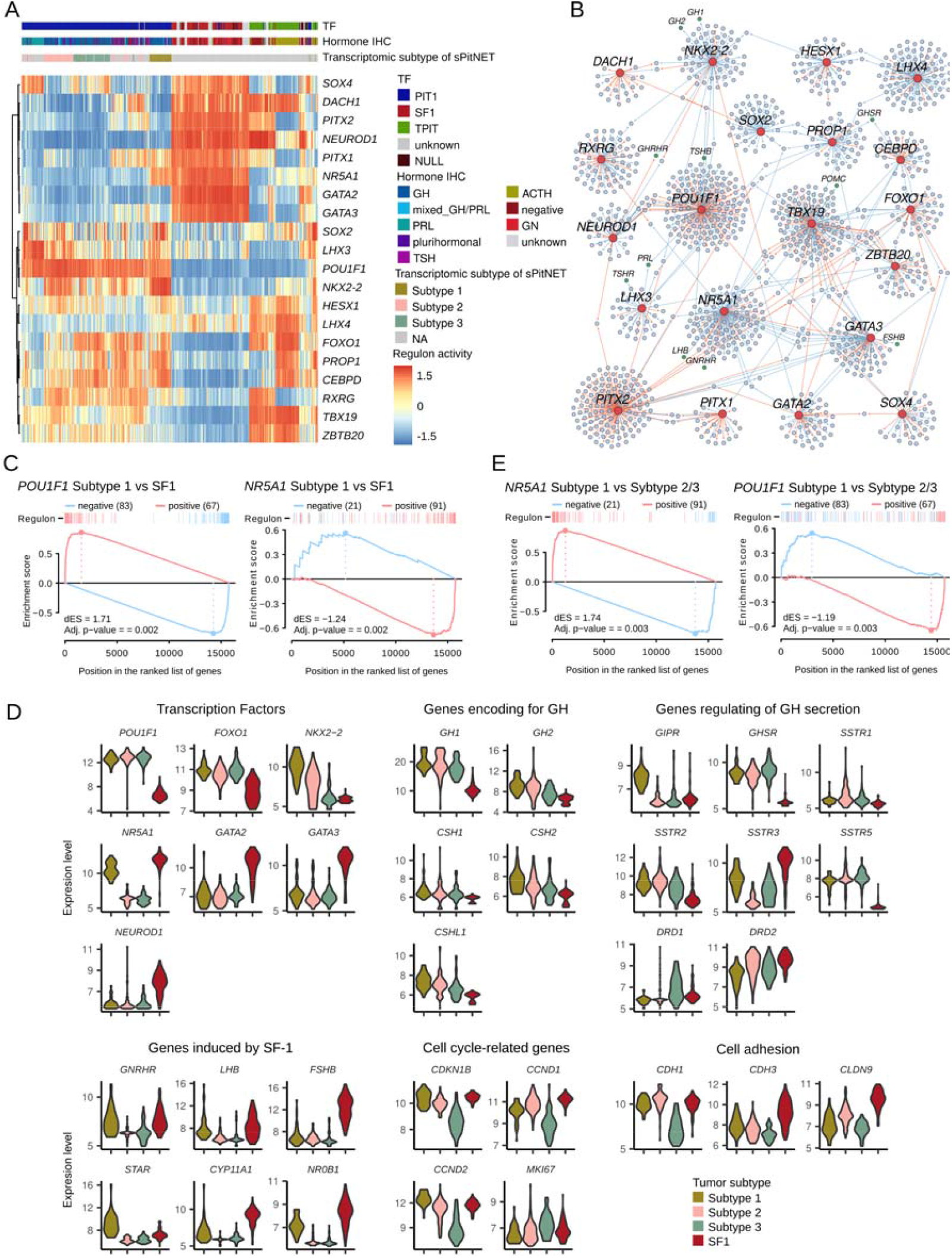
Result of regulon analysis. A) Regulon activity of chosen transcription factors (TF) orchestrating the differentiation of the pituitary secretory cells lineages in PitNET samples with summary of molecular and histological characteristics. B) Network of genes regulated by chosen TF. C) Result of Gene Set Enrichment Analysis (GSEA) showing enrichment of differentially expressed genes (DEGs) between Subtype 1 (PIT-1/SF-1 positive) somatotroph tumors and gonadotroph tumors in genes from regulons of *POU1F1* and *NR5A1*. D) Expression levels of chosen differentially expressed genes. E) Result of GSEA showing enrichment of selected DEGs between Subtype 1 (PIT-1/SF-1 positive) somatotroph tumors and Subtype 1/2 somatotroph tumors in genes from regulons of *POU1F1* and *NR5A1*.

Especially, notable difference in TFs activity was observed in comparison of PIT-1 and SF-1 PitNETs. Double positive acromegaly-related tumors are characterized by a TFs activity pattern characteristic to PIT-1 not SF-1 tumors, with an exception of the activity of *NR5A1*, which is common for double positive and gonadotroph PitNETs. The other TFs specific for gonadotroph tumors such as *GATA2, GATA3* are inactive in double positive PIT-1/SF-1 Subtype 1 somatotroph tumors. The difference in activity of the TFs in each subtype of somatotroph and gonadotroph tumors reflects their distinct expression levels, as well as the difference in expression of the genes involved in secretory activity of pituitary cells (Fig. 2A).

We complemented the regulon analysis with a differential gene expression analysis. Genes differentially expressed (DEGs) between double positive Subtype 1 acromegaly-related tumors and gonadotroph SF-1 tumors were determined (Supplementary Table S4). Next, we analyzed the enrichment of DEGs with the genes from *POU1F1* and *NR5A1* regulons with GSEA. We observed that DEGs are highly enriched in genes regulated by *POU1F1*, but they are also enriched in the regulon of *NR5A1*. Most of the genes which are transcriptionally activated by *NR5A1*, according to the calculated regulon, have lower expression in double positive sPitNETs than in gnadotroph tumors (Fig. 2C). This result indicates that, to some extent, *NR5A1* regulates different set of genes in gonadotroph and Subtype 1-acromegaly tumors (Fig. 2C). Indeed, some known *NR5A1*-depenedent, experimentally validated genes as *CYP11A1, NR0B1*, or *FSHB* are expressed on lower level in double positive somatotroph PitNETs and gonadotroph tumors while the others as *GNRHR* or *LHB* have comparable expression (Fig. 2D). On the other hand, some *NR5A1*-regulated genes as *STAR* have higher expression in Subtype 1 somatotroph than gonadotroph PitNETs.

We also compared Subtype 1 somatotroph tumors to other acromegaly-related tumors (combined Subtype 2 and 3) and looked for the enrichment of DEGs in the regulons of *NR5A1* and *POU1F1* (Supplementary Table S4). DEGs were highly enriched for *NR5A1*-regulated genes but very modestly enriched for the genes regulated by *POU1F1* (Fig. 2E) confirming the specificity of regulation by *NR5A1* to Subtype 1 tumors and showing only slightly different activity patter of *POU1F1*.

We also used the regulon data to determine the TFs that are the most specific to double positive PIT-1/SF-1tumors over the rest of the PitNETs (Supplementary Table S5). This showed *NKX2-2* activity was the most specific (AUC 0.98) confirming the previous suggestion that this TF may serve as reliable marker for this specific transcriptomic subtype [50]. Interestingly, the genes specific to somatotroph cells as *GH1, GH2* and *GHRHR* were annotated with the regulon of *NKX2-2* (Fig. 2B).

### Analysis of scRNAseq data from normal pituitary for identification of normal double positive *NR5A1*/*POU1F1* cells

We analyzed the available single-cell RNAseq data from 10 samples of normal adult pituitary gland (APG). Unsupervised clustering detected 19 clusters, which were classified into 12 cell types based on marker genes and cluster similarity: 4 PIT1 lineage (PIT1.Pro, Somato, Lacto, Thyro), 2 SF1 lineage (Gonado, Gonado.PIT), 1 TPIT lineage (Cortico), 3 immune cells (Macrophages, Monocytes, CD8+ T cells), 1 stem cells (Stem), 2 stroma cells (Endothelial cells, Fibroblasts) (Fig. 3A). Next, we checked whether the identified clusters contain the cells expressing both *POU1F1* and *NR5A1*, and we found such cell to be restricted to clusters of PIT-1 and SF-1 cell lineages. We found a total of 91 *NR5A1/POU1F1* double positive cells (0.6% of all cells) including 49 and 42 in PIT-1 and SF-1 clusters respectively (Fig. 3B). Interestingly, *NR5A1/POU1F1* double positive cells forms a specified subcluster among SF-1 gonadotroph cells, while they are dispersed through the entire cell population PIT-1-positive lineage (Fig. 3C).

**Fig. 3.**
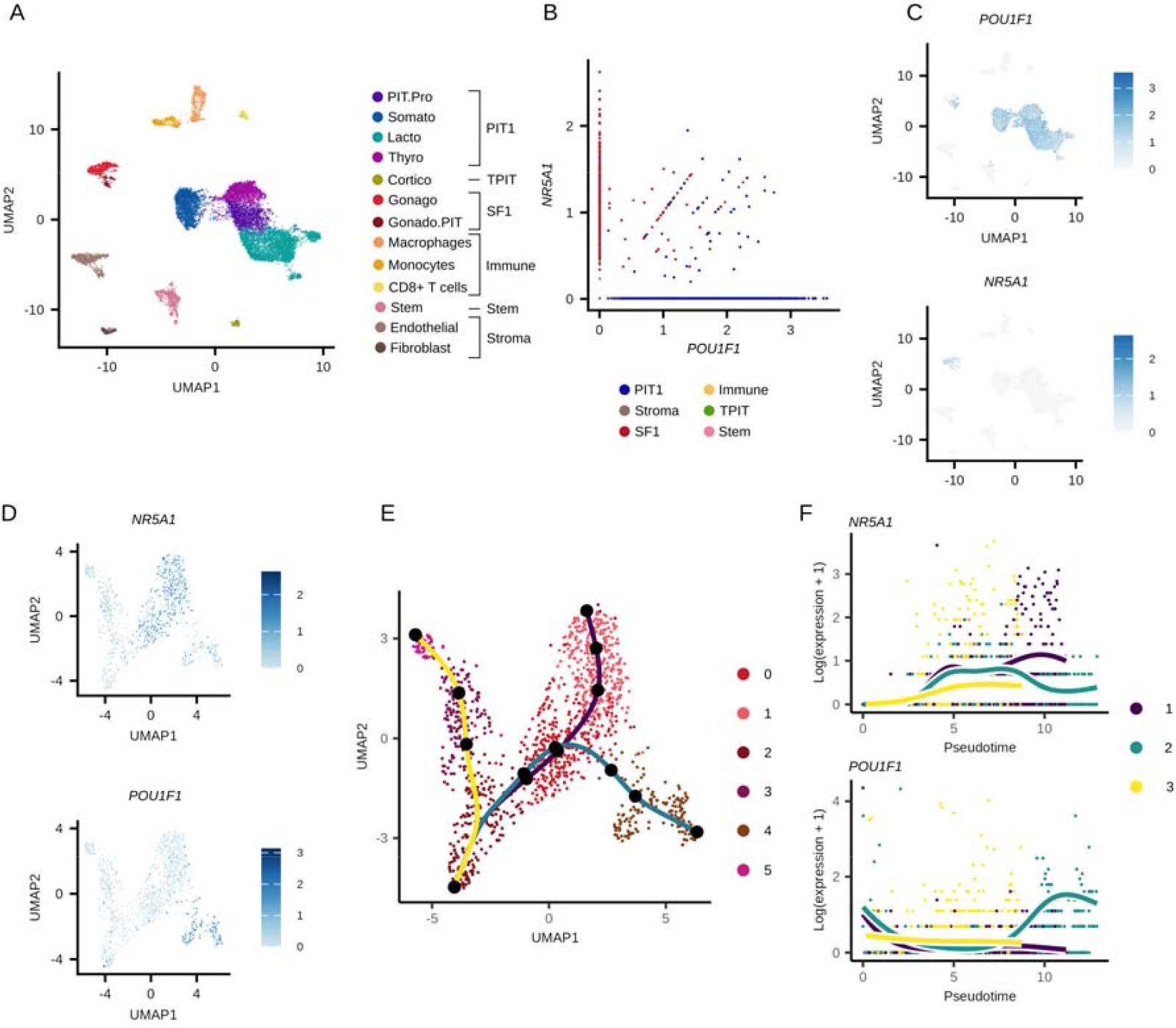
Singel-cell RNAseq data from adult pituitary gland (APG). A) UMAP of cells from APG samples with assigned cell types. B) Co-expression of *POU1F1* and *NR5A1* in cells from APG labeled according to main clusters identified as particular cell subtypes. C) Expression of *POU1F1* and *NR5A1* in cells from APG. D) Expression of *POU1F1* and *NR5A1* in gonadotroph cells from integrated data of APG and fetal pituitary. E) Clustering and differentiation trajectory of gonadotroph cells from integrated data of APG and fetal pituitary. F) *POU1F1* and *NR5A1* expression along differentiation trajectory of gonadotroph cells from APG and fetal pituitary.

Additionally, we verified the presence of such cells in publicly available scRNA data of human fetal pituitary (FPG) as well as mouse and rat pituitary gland. Such double positive cells were observed in human APG (see Figure 3), FPG as well as in mouse and rat (Supplementary Fig. S2). To understand the developmental origin of *NR5A1/POU1F1* double positive cells, we integrated scRNAseq data from human APG and FPG, and processed the data to identify cells from the basic pituitary lineages and performed trajectory analysis focused on cluster of gonadotroph cells. It showed that double positive cells form a subcluster among SF-1 cells (Fig. 3D). They are at the end of separate differentiation trajectory (Fig. 3E). In the cells of this lineage, the expression of *POU1F1* increases, while *NR5A1* decreases with time (Fig. 3F).

### Neoplastic and normal somatotroph cells

Intriguingly, *NR5A1/POU1F1* double positive-cells were identified among both PIT-1 and gonadotroph lineages of normal pituitary gland but RNAseq data from tumors shows that double positive PIT-1/SF-1 tumors are in general similar to PIT-1 positive not gonadotroph PitNETs. To focus specifically on the pituitary tumor cells, we acquired and analyzed the publicly alliable scRNAseq data of somatotroph and gonadotroph PitNETs (25 and 5 samples respectively, Supplementary Table S6). We assigned each sample of somatotroph tumors to one of the previously defined transcriptomic subtypes based on classification of their pseudobulk expression profile. Three tumors were identified as corresponding to Subtype 1 (SF-1/PIT1-positive), 14 as Subtype 2 while 8 as Subtype 3 (Fig. 4A). Then, we integrated scRNAseq data from somatotroph PitNETs with gonadotroph PitNETs (gPitNETs) and APG (total of 40 samples, 142188 cells) and assigned the tumor and non-neoplastic cell populations to five main clusters (Fig. 4B) (Supplementary Fig. S3). Tumor cells of gonadotroph PitNETs were closely similar to normal gonadotroph cell while all the tumor cells of somatotroph tumors, including those PIT-1/SF-1 positive, were similar to normal cells of PIT-1 lineage (Fig. 4C). Importantly, tumor cells of double positive PIT-1/SF-1 somatotroph tumors were definitely unrelated to a subcluster of normal *NR5A1*/*POU1F1* double positive gonadotroph cells (Gonado.PIT1) observed in scRNAseq data on normal adult. They were unambiguously similar to tumor cells of the other somatotroph tumors as well as cells of normal PIT-1 lineage not to gonadotroph tumors cell nor normal gonadotrophs as also reflected by similarity in the expression of key pituitary cell-related genes (Supplementary Fig. S4).

**Fig. 4.**
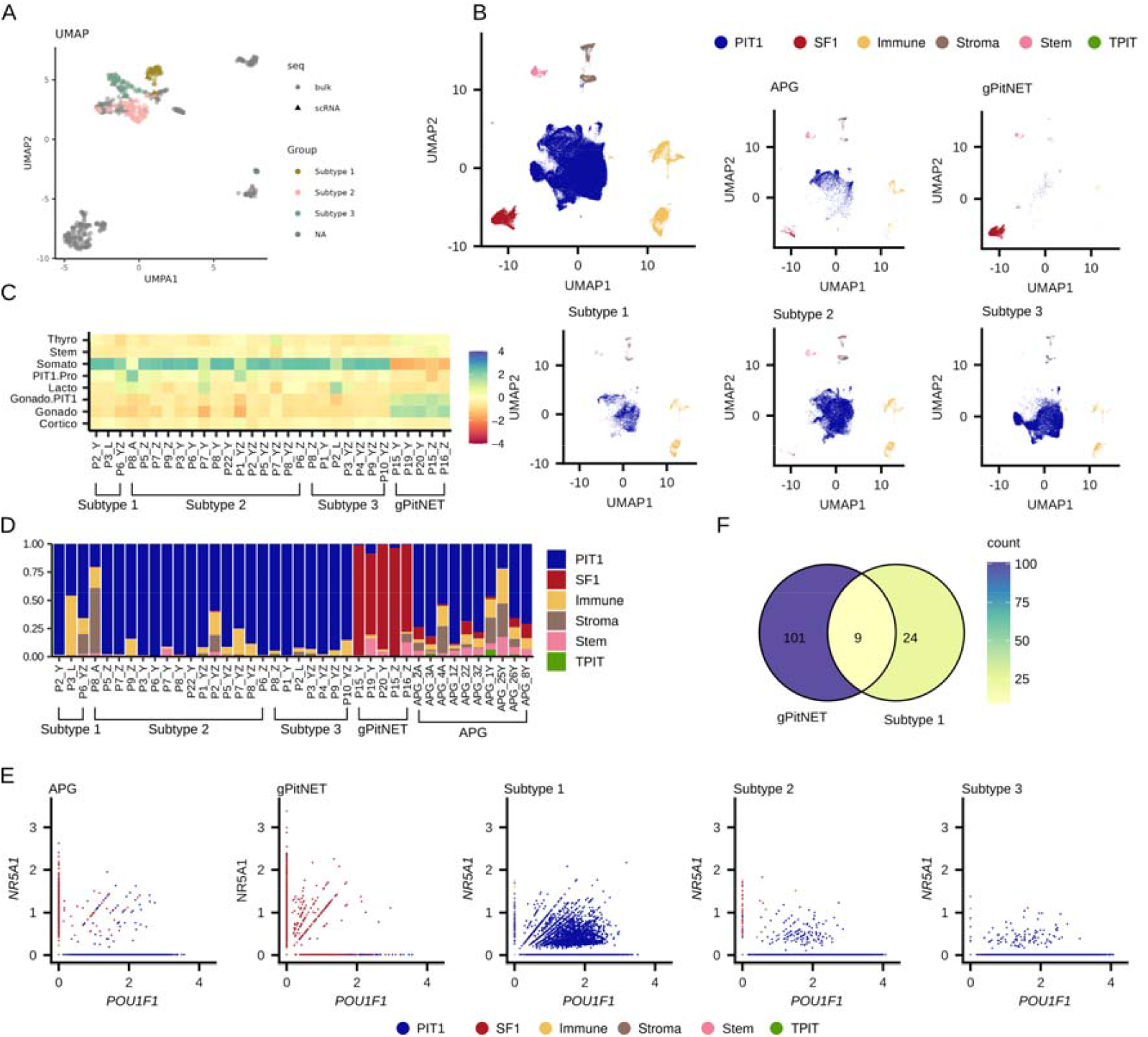
Integrated single-cell RNAseq data of three subtypes of somatotroph PitNETs (sPitNETs), gonadothroph PitNETs (gPitNETs) and adult pituitary gland (APG). A) UMAP of combined pseudobulk counts of samples from scRNAseq of somatotroph PitNETs and bulk RNAseq data of all the somatotroph tumors from study cohort colored by three transcriptomic subtypes of sPitNETs. B) UMAP from integrated data of sPitNETs, gPitNETs and APG with assigned main cell clusters. C) Similarity of cells in specific samples to cell clusters from APG. D) Main cell clusters representation in samples. E) Co-expression of *POU1F1* and *NR5A1* with assigned main clusters in specific sample groups. F) Comparison of overlap in genes from *NR5A1* regulon calculated based on gonadotroph cells from gPitNETs and somatotroph cells from Subtype 1 (PIT-1/SF-1 positive) sPitNETs.

The gonadotroph tumors were basically composed of cells of gonadotroph nature, while each of the subtype of somatotroph tumors included mostly cells of PIT-1 lineage (Fig. 4D). Some low amount of the cells expressing both *NR5A1* and *POU1F1* was detected in gonadotroph tumors (these cells belonged to cluster of SF-1 gonadotroph cells) as wells as in two subtypes of somatotroph tumor subtypes without NR5A1 expression (these cells belonged to cluster of PIT-1 cluster) (Fig. 4E).

We also looked for the abundance of nonneoplastic cells as immune, stromal and stem-like cells in individual tumor samples and APG. A notable intertumoral heterogeneous was observed. A high content of immune cells was identified in 1 of 3 samples of double positive (SF-1/PIT-1) somatotroph PitNETs, that possibly affect the analysis (Fig. 4D). In general, no difference in the frequency of subtypes of nonneoplastic cells between somatotroph tumor subtypes, were found. Lower content of stemm-like cells and monocytes was found in somatotroph tumors in general as compared to APG as detailed in Supplementary Fig. S5.

To verify the observation that a distinct set of genes can be regulated by SF-1 in double positive somatotroph and gonadotroph PitNETs we established regulons of *NR5A1* in gonadotroph tumor cells and double positive somatotroph tumor cell independently. Analyzes of genes in regulons confirmed results from comparison of bulk samples of Subtype 1 and gonadotrhop tumors that *NR5A1* regulates different genes in those PitNET types (Fig. 4F, Supplementary Table S7).

Determining abnormal genes expression in somatotroph tumors by comparison with normal pituitary somatotroph is unavailable with bulkRNAseq due to a complexed cellular composition of pituitary gland. This can be in turn achieved with scRNAseq data. We compared genes expression in tumor cells of each subtype of somatotroph PitNETs with normal somatotrophs of APG. We identified 1149, 1853 and 2021 DEGs for Subtype 1, 2 and 3 respectively, with a set of DEGs unique for each subtype (Fig.5A, Supplementary Table S8). The result confirmed a difference in the expression of important genes related to pituitary functioning and pathogenesis between subtypes, as well as differences between tumor and normal cells. Showing abnormal expression of TFs (*POU1F1, NR5A1*) as wells genes encoding for GH (*GH1, CSH1, CSHL1*), in genes regulating GH secretion (*GHRHR, GIPR, GHSR, SSTRs* and *DRDs*) (Fig. 5B). Results confirmed abnormal increase of the expression of genes induced by SF-1 in Subtype 1 somatotroph tumors (PIT-1/SF-1 positive) as steroidogenesis-related genes and *GNRHR*, but not in *LHB* and *FSHB*. The results also confirmed a tumor-specific decrease of the expression of cell adhesion and cell cycle-related genes in Subtype 3 tumors.

**Fig. 5.**
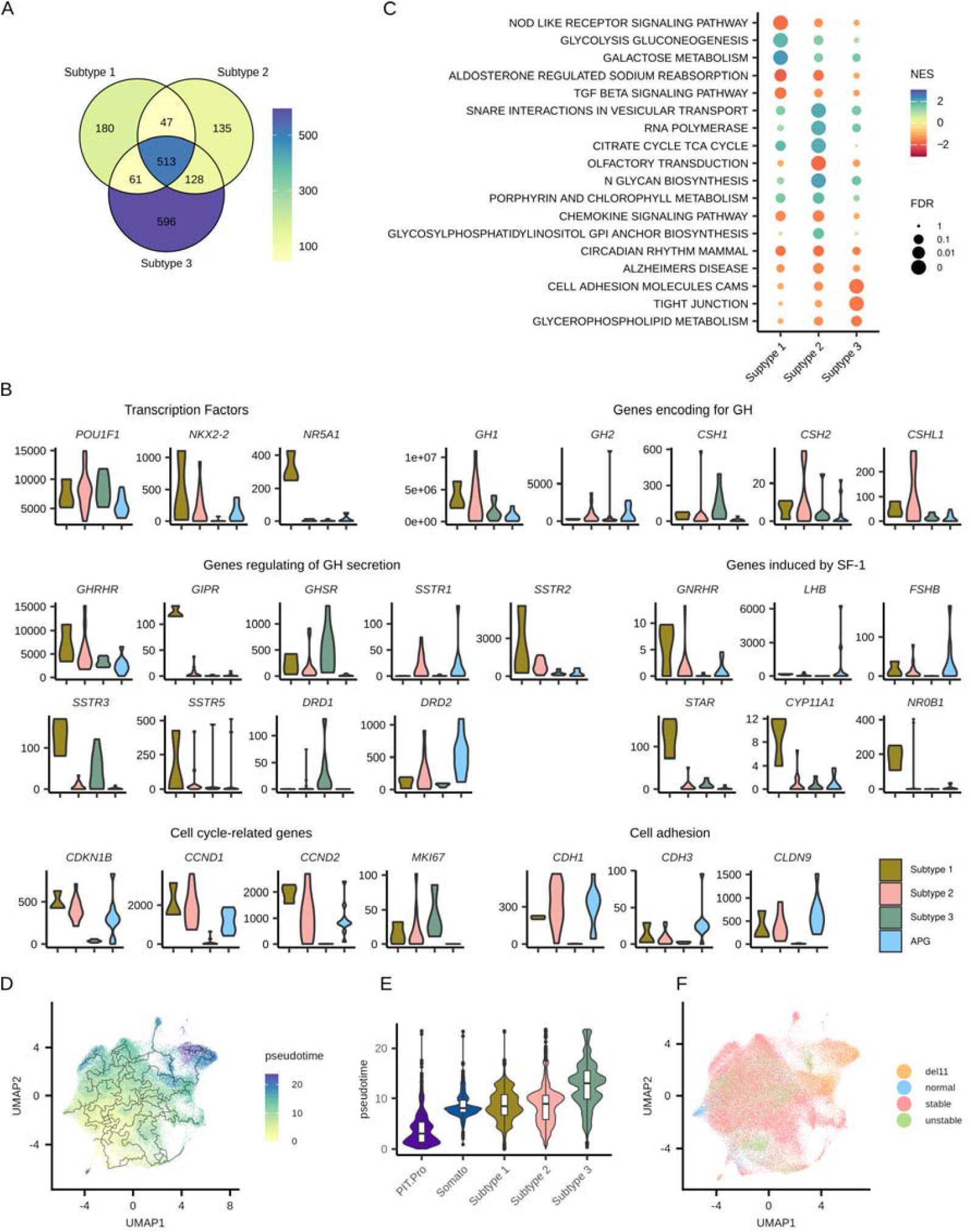
Comparison of tumor somatotroph cells of each particular transcriptomic subtypes of somatotroph PitNETs (sPitNETs) and normal somatotroph cells of adult pituitary gland (APG). A) Overlap of differentially expressed genes (DEGs) identified in comparison of tumor somatotroph cell of each transcriptomic subtype of sPitNETs and normal somatotrophs of APG. B) The expression of selected DEGs in each transcriptomic subtype of sPitNETs and normal somatotrophs. C) Result of gene set enrichment analysis (GSEA) of DEGs from comparison of cells from specific transcriptomic subtypes of sPitNETs to APG based on KEGG database. D) UMAP of somatotroph cells from integrated APG and sPitNET data colored by Monocle3-inferred pseudotime. E) Comparison of distribution of pseudotime in cells from specific cell groups. F) UMAP of somatotroph cells from integrated APG and sPitNET data, classified according to genomic instability into stable, unstable and chromosome 11 deleted.

KEGG pathways enriched for a DEGs found in the comparison of neoplastic cells of each tumor subtype and normal somatotropes were identified (Supplementary Table S9) including the pathways specific for distinct subtypes (Fig. 5C). Aberrant genes expression influence upregulation of galactose and glucose metabolism and downregulation of TGF beta signaling in Subtype 1 tumors; upregulaiton of TCA cycle, in Subtype 2 and downregulation of cell adhesion in Subtype 3.

ScRNAseq data were also used to calculate psudotime-related trajectory of the PIT.Pro and somatotroph cells in APG and somatotroph tumor cells to trace the differences between tumor samples of known subtypes (Fig. 5D). We observed a notable variation of pseudotime among cells from specific Subtypes without a clear relationship to development of normal pituitary cells, however we noted the individual samples of Subtype 3 were the most distant as compared to normal somatotroph (Fig. 5E). Determining copy number profiles with scRNAseq data showed loss of chromosome 11 in these samples (Fig. 5F, Supplementary Fig. S5). This chromosomal loss was already identified as recurrent CNV in subtype 3 tumors [4,6].

## Discussion

The principles of the classification of pituitary tumors is determining from which developmental pituitary lineage they originate [51]. Basically, the lineage of corticotroph cells is characterized by the expression of TPIT transcription factor, gonadotroph cells express SF-1 TF and PIT-1-positive lineage includes somatotropes, lactotropes and tyreotropes. In general, PitNETs falls in the three basic categories of TPIT, SF-1 and PIT-1 tumors and the tumors developed from distinct pituitary cell lineages have distinct molecular profile as observed in transcriptomic, epigenetic and proteomic levels [15,19,21]. PitNETs of a particular type are still characterized by a degree of heterogeneity in clinical, histological and molecular features.

Clinically functioning somatotroph pituitary tumors fall into 3 distinct molecular subtypes which are not clearly distinguished in official WHO classification [4–7] and one of these subtypes is characterized by the expression of *NR5A1* (SF-1) TF. Such double positive PIT-1 and SF-1-expressing sPitNETs were also occasionally, previously reported by others [9–15] and they were pointed as a matter of interest. Preliminary goal of our study was to identify the tumors of this subtype in each publicly available RNAseq data set from sPitNETs. Indeed, using unsupervised transcriptomic analysis of 193 tumor samples, we found that ∼20% of sPitNETs are double positive tumors, confirming their previously observed frequency among acromegaly-related tumors. Importantly, they were detected in each of 5 data sets of various patients’ population (Asian, European, South American) included in the analysis.

The developmental identity of tumors co-expressing PIT-1 and SF-1 has been recently discussed in the literature, and they were differently referred by the authors as multilineage pituitary tumors [16,17,50,52,53], gonado/somatotroph tumors [6], somatogonadotroph [18] or plurihormonal tumors [9,11,54]. We addressed the question whether these acromegaly-related sPitNETs truly have molecular features of pituitary gonadotropes and to what extent they resemble SF-1 tumors. The results of our analysis comprising 546 pituitary tumor samples showed that, in terms of genes expression, double positive sPitNETs are closely similar to the other PIT-1 not to gonadotroph tumors. By the use of scRNAseq data, we were able to compare tumor cells of each subtype of somatotroph tumors and gonadotroph PitNETs with normal pituitary cells of both PIT-1 and SF-1 lineages. It clearly confirmed that double positive Subtype 1 sPitNETs are counterparts of normal PIT-1 cells, not pituitary gonadotropes.

Results of comparing the activity of known TFs involved in differentiation of pituitary cell lineages is in line with the above observation. PIT-1/SF-1 sPitNETs are characterized by activity of TFs known to be involved in development of PIT-1, not SF-1 lineage as *POU1F1, PROP1, CEBPD* or *FOXO1* [55]. Beside the activity of SF-1 (*NR5A1*), these tumors do not show the activity of the other known gonadotroph lineage-related TFs as *GATA2* or *GATA3* [56].

Part of PIT-1/SF-1 positive sPitNETs were found to express LH in immunohistichemical staining [6,7,18]. Our analysis confirms that these tumors indeed express both *LHB* and *GNRHR* at the level similar to gonadotroph tumors. Both these genes are regulated by SF-1 transcription factors in pituitary gonadotroph cells [57,58]. Interestingly, a category of somatotroph pituitary tumors that responds with increased GH secretion to luteinizing hormone-releasing hormone (LHRH) was recently described [59]. Most of these tumors also showed paradoxical increase of GH in response to oral glucose tolerance test (OGTT) [59] a phenomenon observed in a specified group of patients with acromegaly [60] as a result of ectopic expression of *GIPR* [61]. *GIPR* expression is correlated with *NR5A1* level in sPitNETs [7] and it corresponds to demethylation at *GIPR* promoter that was observed in SF-1/PIT-1 positive sPitNETs [5]. The presented analysis of both bulk- and scRNAseq confirms that these double positive sPitNETs are those that express high level of *GIPR* and expression of this gene distinguishes the tumor cells of these subtype from normal pituitary somatotropes, which are GIPR-negative. These tumors also express *LHB* and *GNRHR*, therefore we conclude that Subtype 1 sPitNETs are those that increase GH secretions after LHRH or glucose load.

Since the expression of *LHB* and *GNRHR* is related to *NR5A1* in both Subtype 1 sPitNETs and gPitNETs the other genes regulate by this TF turn out to be differentially expressed in these tumor types. The best validated gene positively regulated by SF-1 is *STAR*, which encodes a protein playing a key role in steroidogenesis in adrenal cortex and ovary [62]. We clearly observed high expression of *STAR* in Subtype 1 sPitNETs, but not in gonadotorph tumors. In opposite, the other SF-1 regulated genes such as *FSHB, CYP11A1* and *NR0B1* (DAX1) have higher expression level in gonadopoph tumors. More general, our analysis of *NR5A1* regulons in bulk- and scRNAseq data indicate that this TF regulates somewhat different set of genes in double positive sPitNETs and gonadotropinomas. Our regulon analysis also confirms the previous report that *NKX2-2* transcription factor can be considered as specific to PIT-1/SF-1 sPitNETs [50] as we observed the highest activity of *NKX2-2* in this tumor subtype.

The analysis scRNAseq from normal APG showed the presence of a few PIT-1/SF-1 cell among both PIT-1 and gonadotroph lineage cells. Similarly, such cells were identified in human fetal pituitary as well as in pituitary from mouse and rat. Interestingly, normal PIT-1/SF-1 double positive cells represented a specified developmental subcluster of gonadotroph cells, but double positive tumor cells from Subtype 1 sPitNETs turn out to be completely dissimilar (unrelated) to this gonadotroph cluster. Thus, we conclude that these tumor cells originate from PIT-1 lineage not from gonadotroph cell subtype.

Taking advantage of the scRNAseq data, we compared genes expression in each subtype of sPitNETs with normal somatotrophs to identify tumor-specific aberrantly expressed gens. The profile of aberrant genes expression in tumor cells of each sPitNET subtypes confirms the observation of difference in genes expression, both observed in current bulk RNAseq results and previously reported [6,7]. Additionally, by integrating the scRNAseq data from normal pituitary and sPitNETs we made an attempt of pseudotime analysis to match tumor samples with normal developmental trajectory from ProPit cell to somtotroph lineage. The result shows that sPitNETs are more related to mature somtotroph lineage and the clear difference in development trajectory of tumor of three subtypes could not be found. The interesting results from this analysis was that Subtype 3 tumors are the least related to normal somatotrophs and these are the tumors characterized by chromosome 11 loss. This cytogenetic change was already recognized as recurrent DNA copy number variant in sparsely granulated sPitNETs [4,6].

In this study, we were able to collect a large data set of bulk RNAseq results from PitNETs, but much lower number of scRNAseq data are currently available. Combining the data from 5 studies we identified only 3 PIT-1/SF-1 sPitNETs what should be considered as main limitation of our study. Low number of these sample influence the statistical inference power, while variability within this group prompt us to cautiously interpret the single cell data. In example, the results indicate the increase of *SSTR2* in this tumor subtype as also reported previously [16], but this observation is notably biased by very high expression of this SSTR in 1/3 samples. The role of immune component of tumors environment in sPitNEt has been described [63] and scRNAseq results allows for an in depth analysis of the microenvironment cell composition. Unfortunately, given intertumoral heterogeneity, we were not able to draw a clear conclusion on difference in microenvironment difference in three subtypes of sPitNETs.

## Conclusions

From the perspective of transcriptomic profiling, we do not find the rationale for considering PIT-1/SF-1 sPitNETs as multilineage or “somatogonadotoph” tumors. They have genes expression pattern and profile of the activity of TFs very similar to the other somatotroph tumors with no similarity to gonadotroph tumors with the exception of *NR5A1* expression and expression of some of SF-1-regulated genes as *LHB* and *GNHRH*. SF-1 apparently regulates slightly different set of genes in these sPitNETs and gonadotroph tumors, with different *STAR* expression as an example. Accordingly, tumor cells of PIT-1/SF-1 sPitNETs are closely related to normal pituitary somatotrophs not gonadotrophs, however a subtype of normal gonadotroph cells with *POU1F1* expression was also identified.

## Declarations

### Availability of data and material

Freely available datasets from bulkRNAseq (accession numbers: HRA003588, E-MTAB-11889, E-MTAB-7768, GSE213527, GSE209903) and scRNAseq (accession numbers: PRJCA009690, GSE208108, HRA003110, HRA003483, PRJNA1137596 and GSE142653).

### Authors’ contributions

Author Contributions: Conceptualization, JR, MB; Methodology, JR, MB, QZ; Formal analysis JR, MB, QZ; investigation, JR, MB, QZ; resources, JR, MB, QZ; data curation, JR, MB; writing— original draft preparation, JR, MB; writing—review and editing, all the authors; visualization, JR ; supervision, MB; project administration MB; funding acquisition, MB.

### Funding

This research was funded by Maria Sklodowska-Curie National Institute of Oncology, grant SN/GW5/2023.

### Ethics approval and consent to participate

not applicable; secondary use of data

### Consent for publication

Not applicable

### Competing interests

The authors declare no conflict of interest.

## Acknowledgements

Not applicable

## Notes

### Competing Interest Statement

The authors have declared no competing interest.

