## Supplementary Fig. for "Transcriptomic perspective on acromegaly- related PIT-1/SF-1 positive pituitary tumors"

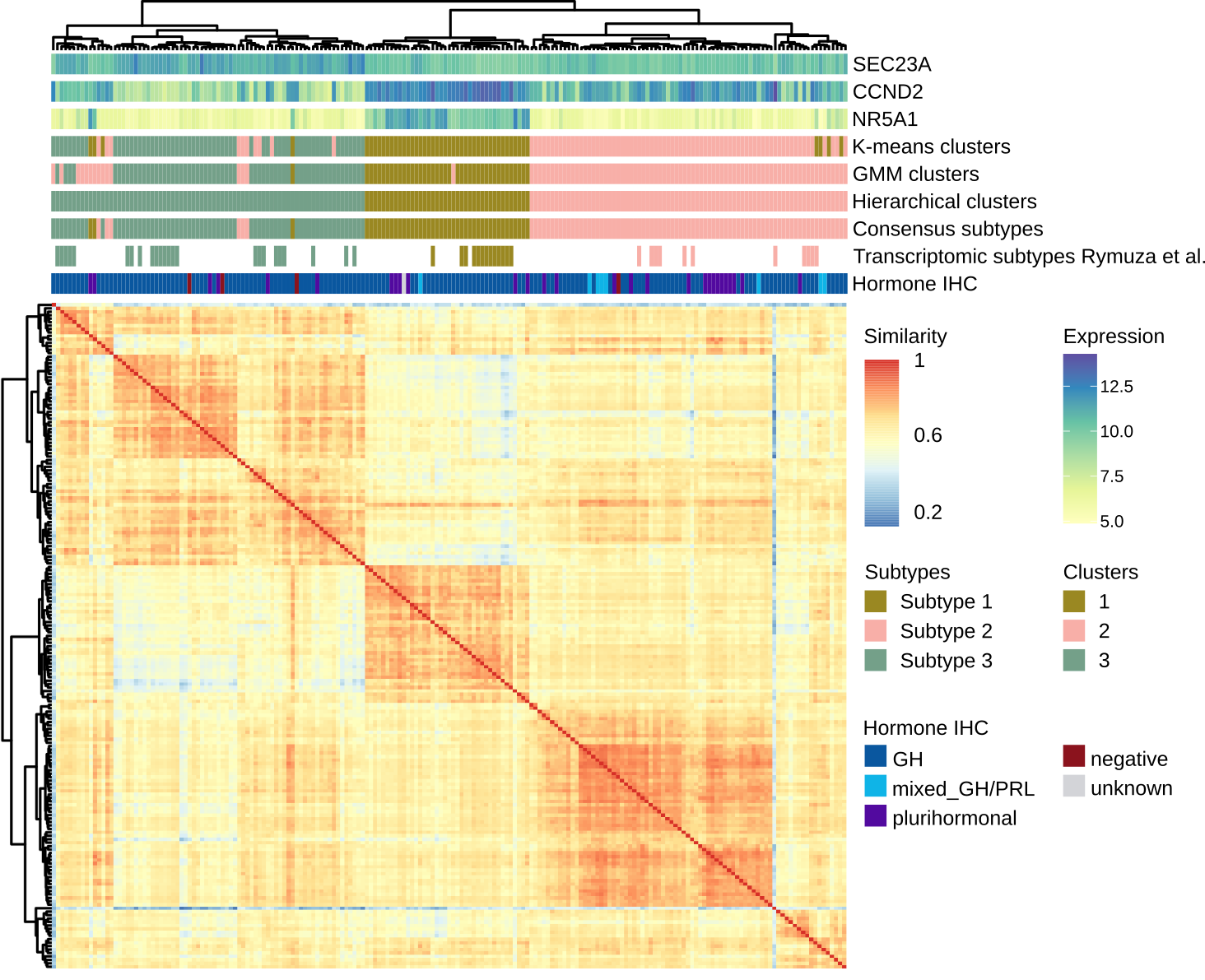

Fig. S1: Clustering of acromegaly-related tumors with results of different clustering algorithms, expression of subtype marker genes, histological groups and sample similarity matrix.

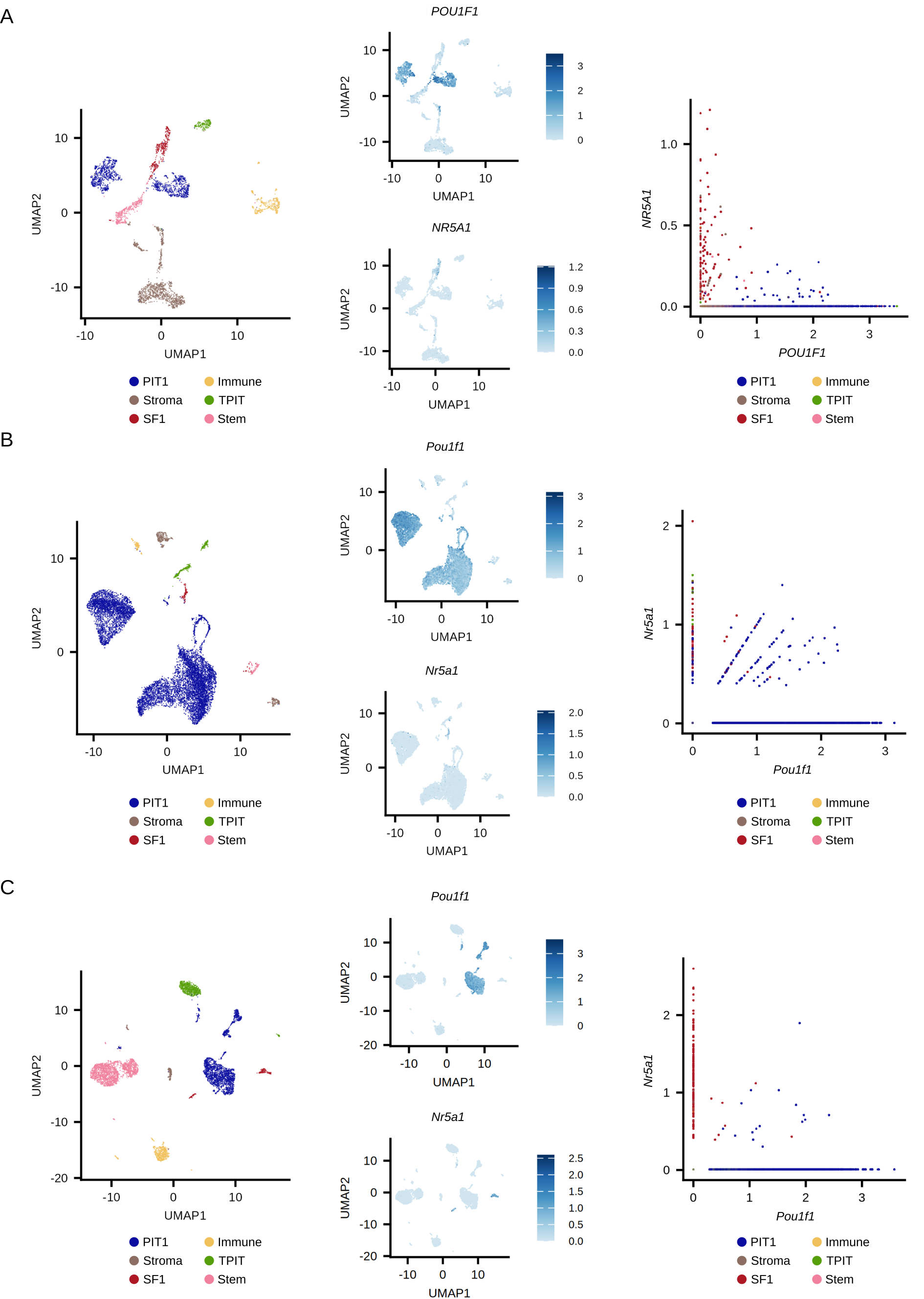
Fig. S2 UMAP with annotated main cell clusters, expression of *POU1F1 (Pou1f1)* and *NR5A1 (Nr5a1)*, co-expression of *POU1F1 (Pou1f1)* and *NR5A1 (Nr5a1)* from cells from fetal pituitary gland (A), mouse pituitary gland (B), rat pituitary gland (C).

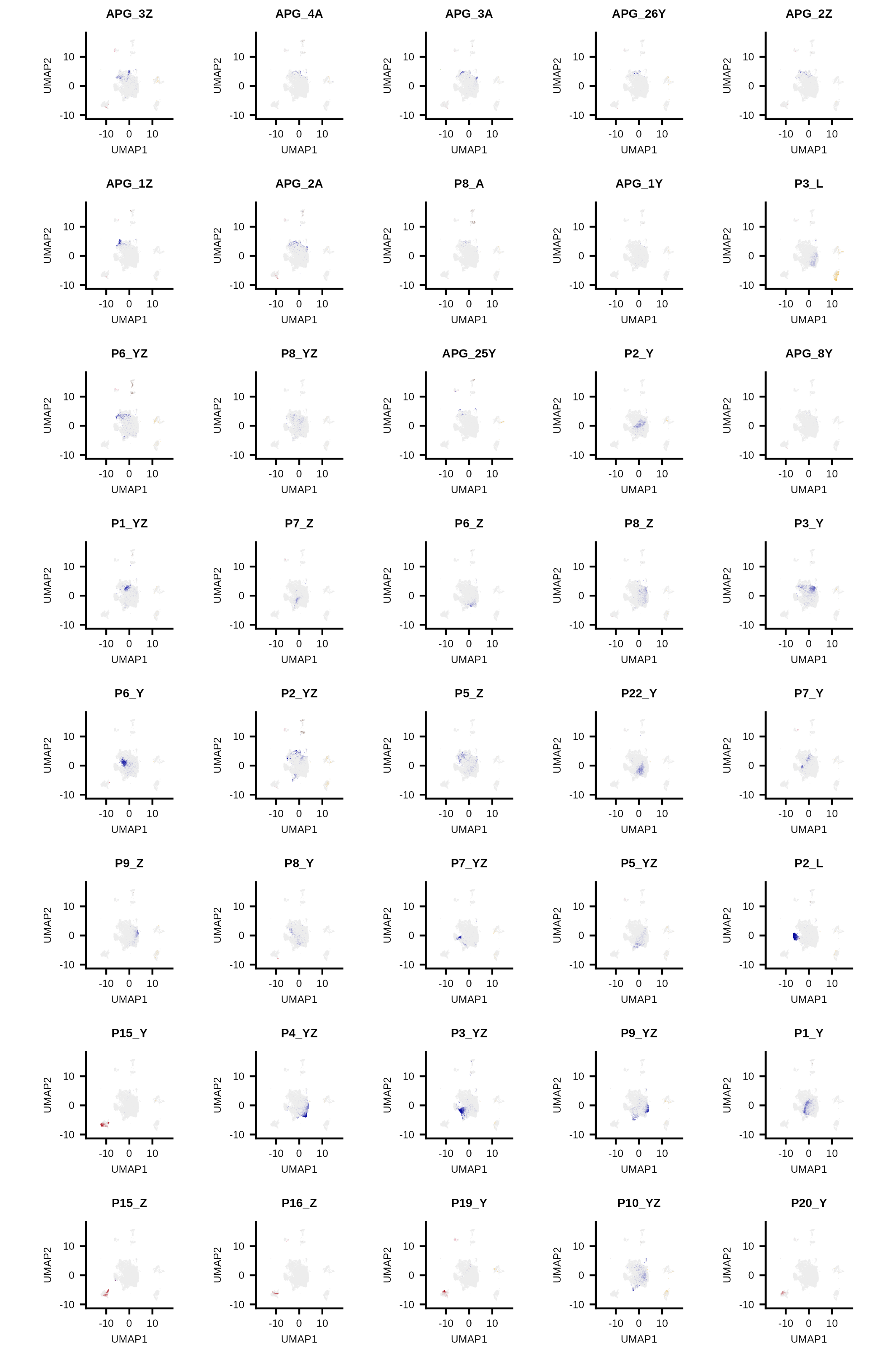
Fig. S3 UMAP from integrated data from somatotroph PitNETs, gonadothroph PitNETs and adult pituitary gland with highlighted cells from specific sample.

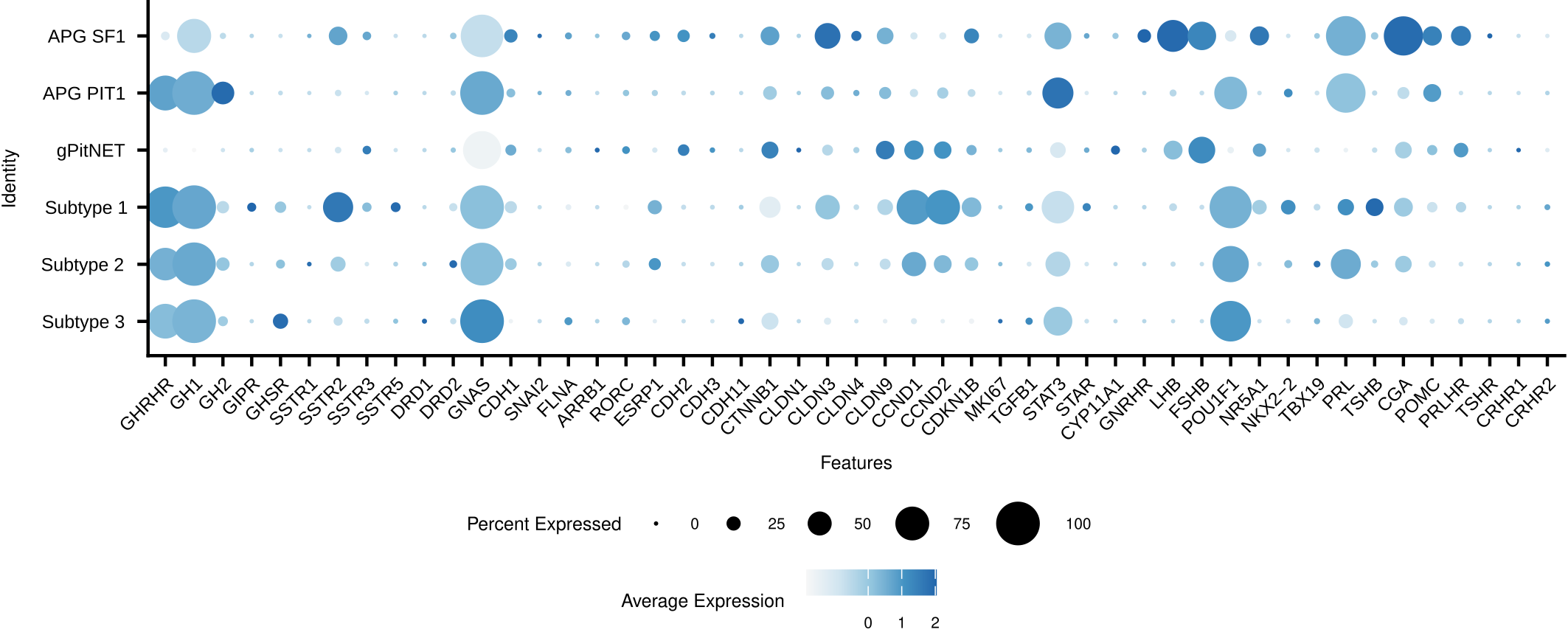
Fig. S3 Expression of key pituitary cell-related genes in different cell groups.

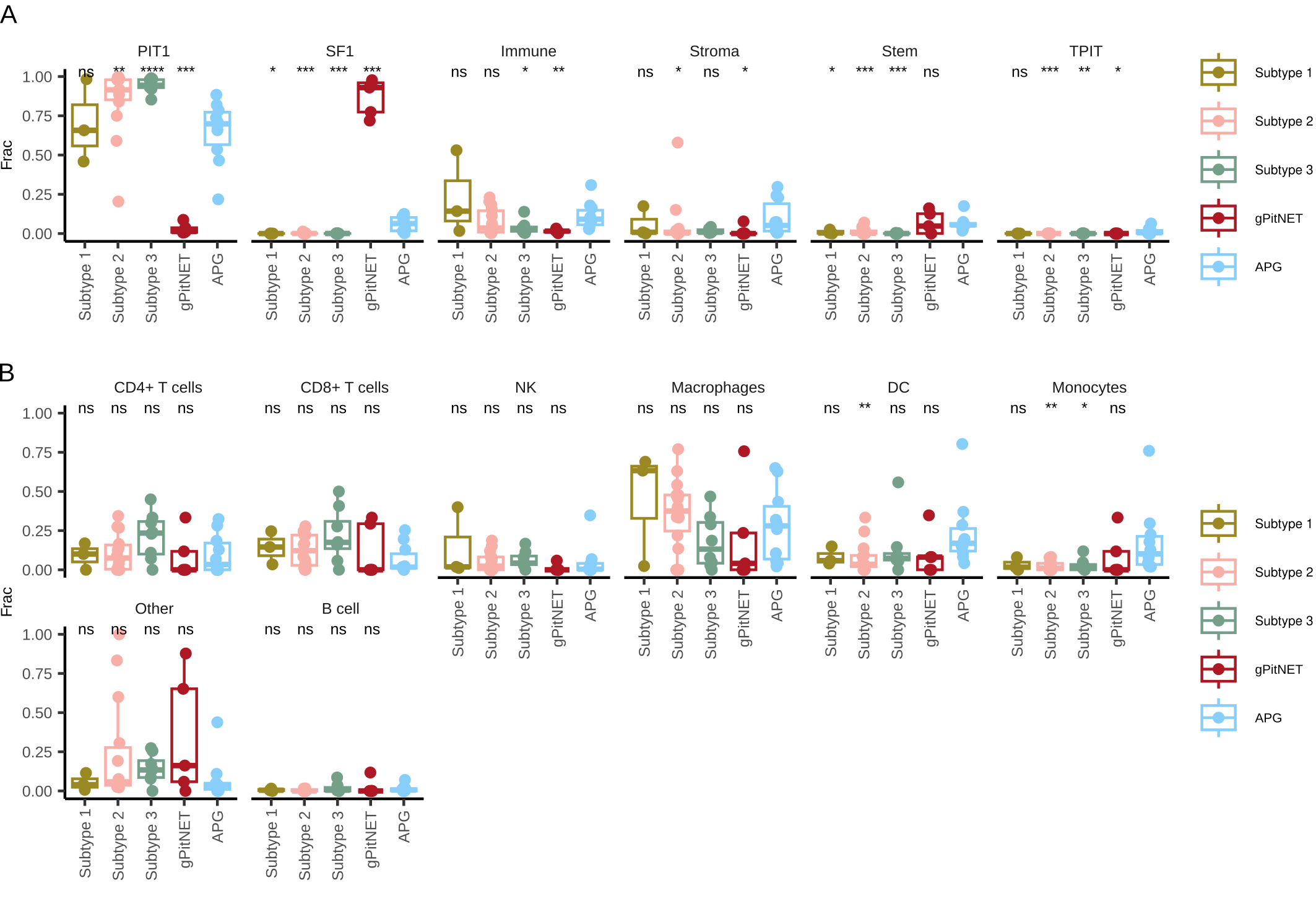

Fig. S4 Abundance of different cell groups in sample groups at the level of main groups (A) and different immune cells (B).

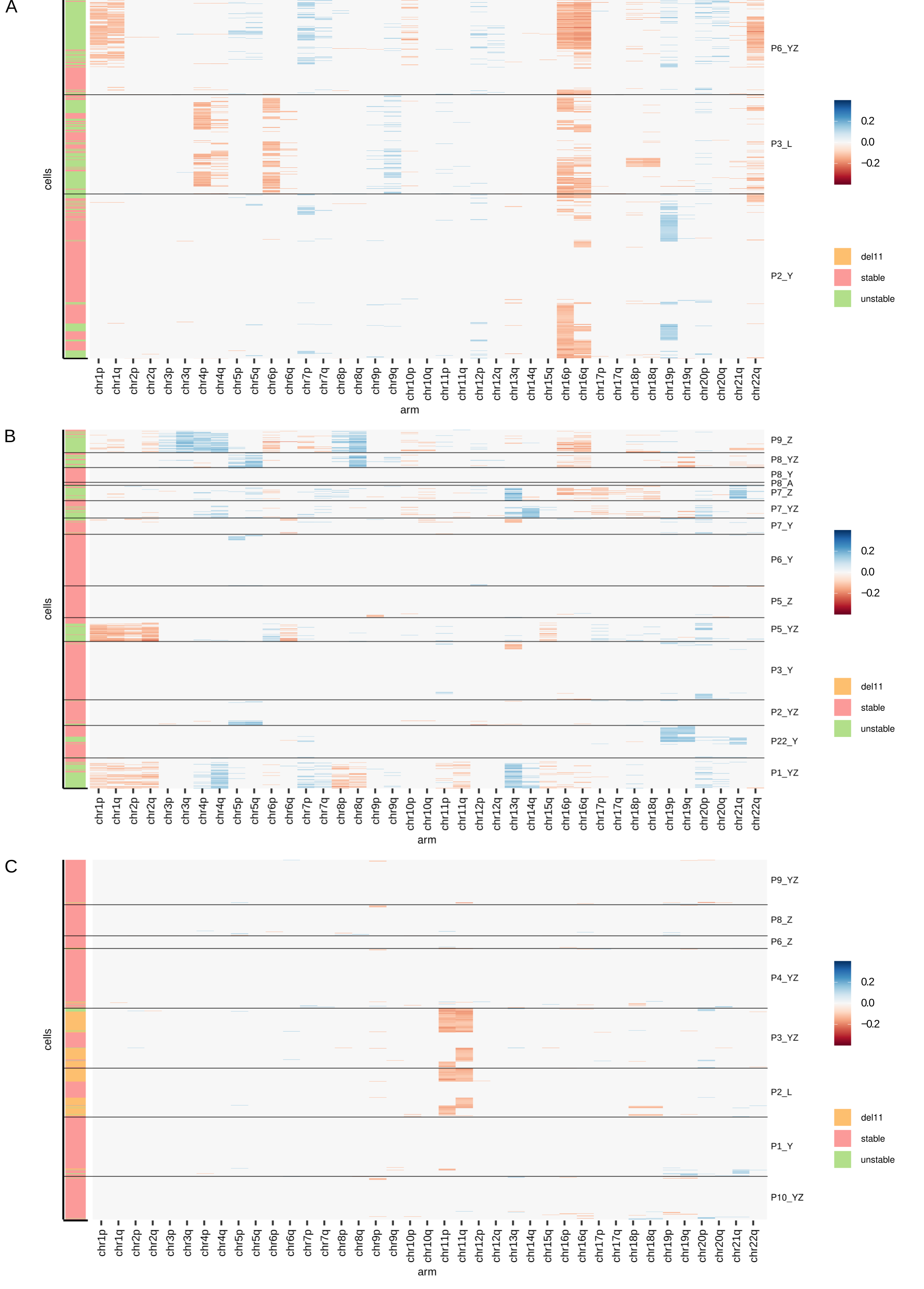
Fig. S5 Heatmap of inferred copy number profiles with cells assigned to stability clusters of cells from Subtype 1 (A), Subtype 2 (B), Subtype 3 (C).
